# Strong information-limiting correlations in early visual areas

**DOI:** 10.1101/842724

**Authors:** Jorrit S Montijn, Rex G Liu, Amir Aschner, Adam Kohn, Peter E Latham, Alexandre Pouget

**Author notes:** These authors contributed equally (Jorrit S Montijn, Rex G Liu).

## Abstract

If the brain processes incoming data efficiently, information should degrade little between early and later neural processing stages, and so information in early stages should match behavioral performance. For instance, if there is enough information in a visual cortical area to determine the orientation of a grating to within 1 degree, and the code is simple enough to be read out by downstream circuits, then animals should be able to achieve that performance behaviourally. Despite over 30 years of research, it is still not known how efficient the brain is. For tasks involving a large number of neurons, the amount of information encoded by neural circuits is limited by differential correlations. Therefore, determining how much information is encoded requires quantifying the strength of differential correlations. Detecting them, however, is difficult. We report here a new method, which requires on the order of 100s of neurons and trials. This method relies on computing the alignment of the neural stimulus encoding direction, **f′**, with the eigenvectors of the noise covariance matrix, **Σ**. In the presence of strong differential correlations, **f′** must be spanned by a small number of the eigenvectors with largest eigenvalues. Using simulations with a leaky-integrate-and-fire neuron model of the LGN-V1 circuit, we confirmed that this method can indeed detect differential correlations consistent with those that would limit orientation discrimination thresholds to 0.5-3 degrees. We applied this technique to V1 recordings in awake monkeys and found signatures of differential correlations, consistent with a discrimination threshold of 0.47-1.20 degrees, which is not far from typical discrimination thresholds (1-2 deg). These results suggest that, at least in macaque monkeys, V1 contains about as much information as is seen in behaviour, implying that downstream circuits are efficient at extracting the information available in V1.

## Introduction

A fundamental question in neuroscience is: do neural circuits, and in particular cortical circuits, perform efficient computation over behaviourally relevant variables? If computation is efficient, then behavioural performance should be only slightly worse than neural performance; if it is inefficient, it should be much worse. For example, humans have an orientation discrimination threshold of about 1 degree (Orban, Vandenbussche, and Vogels 1984; Webster, Switkes, and De Valois 1990; Mäkelä, Whitaker, and Rovamo 1993; Mareschal and Shapley 2004). If an ideal observer of V1 activity can read out orientation with an error of about 1 degree, then computation (of orientation) is efficient; if an ideal observer has an error of 0.001 degrees, then computation is highly inefficient. The answer to this question is important because it gives insight into computational strategies.

Despite 30 years of research, we still do not know the answer to this question. According to Shadlen et al (1996), the population response in area MT contains roughly the same information as observed in the behaviour for a simple motion discrimination task. This result seems surprising because there are many neurons in area MT whose individual responses convey only slightly less information than is available in the behaviour (Shadlen and Newsome 1996). Given that thousands of MT neurons encode direction of motion, the population information could potentially be much larger than the behavioural information. However, Shadlen et al (1996) suggested that the population information is in fact small due to the presence of large pairwise noise correlations among neurons with similar tuning, with correlation coefficients around 0.2 on average. As a result, information saturates as the number of neurons increases, and does so at a value close the one seen in behaviour (Saturation Low **FIG 1**).

**Figure 1.**
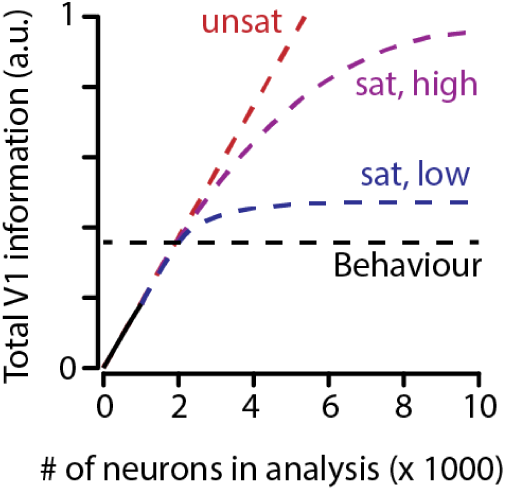
Information in neural circuits, and across brain areas. Information in early sensory areas can scale in different ways with the number of neurons. According to Shadlen et al (1996), information in MT saturates to a value close to the behavioural information (black horizontal line), suggesting that downstream cortical circuits are efficient (blue curve). On the other extreme, information might scale linearly with the number of neurons (red curve), in which case cortical circuits downstream would be particularly inefficient, since a small fraction of that information would make it to the behaviour. In an intermediate scenario, information might saturate at a finite value, but still much larger than what is seen in behaviour. This would again correspond to inefficient cortical circuits, since much of the information would be lost.

This conclusion, however, has been challenged by subsequent theoretical studies. For a population of neurons with heterogeneous tuning curves that are approximately random with respect to the noise covariance matrix, information is proportional to the size of the population (Shamir and Sompolinsky 2006; Ecker et al. 2011) (Unsaturated, **FIG 1**). Of course, the constant of proportionality could be very small, so that with realistic numbers of neurons there would not be much information in the neural population. However, this possibility is inconsistent with the observation that single neurons carry information that is close to the behavioural information in MT (Britten et al. 1996), V4 (Cohen and Maunsell 2010) and V1 (Graf et al. 2011). Thus, if information is linear in the number of neurons, the slope must be sufficiently large that the information in even a small number of neurons would be much larger than what is suggested by behaviour.

The problem with these theoretical studies is that they didn’t take into account the fact that information-limiting correlations, also known as differential correlations (Moreno-Bote et al. 2014), must be large. Differential correlations correspond to fluctuations in population activity that line up with the vector of tuning curve derivatives, denoted **f′**. If the variance of these fluctuations is proportional to the number of neurons, information saturates (Moreno-Bote et al. 2014). Detecting such correlations experimentally is hard because, while being large, they tend to be small compared to non-information-saturating correlations. Nonetheless, for sufficiently large populations, such correlations must exist in cortex. That’s because there is only a finite amount of information entering the brain, which precludes the possibility of information increasing forever as the number of neurons increases. The critical questions, then, are: how big are these information limiting correlations, and at what level does information saturate relative to behavioural information (Saturation Low versus High, **FIG 1**)?

In principle, this debate could be settled with a conceptually simple experiment: record the spiking activity of many neurons simultaneously, measure information as a function of the number of neurons, and ask whether it saturates near the behavioural information. Unfortunately, it is often difficult to detect information saturation in populations of ∼100 neurons, the population size typically available in a single area from electrophysiological recordings (Semedo et al. 2019; Jun et al. 2017). Calcium imaging can provide recordings from 1000s of neurons (Ahrens et al. 2013; Stringer et al. 2019), but it is subject to a variety of problems: the calcium signal has a long time constant, making it difficult to determine activity in short time windows, and extracting spikes is imperfect, making the signal noisy. It is therefore important to have methods that can infer information saturation from the spiking responses of 100s of neurons

Here we present such a method. It takes advantage of the fact that a population of neurons in which information saturates must contain large differential correlations. As we will show below, the presence of large differential correlations leads to a characteristic, and detectable, signature in the eigenspectrum of the population’s covariance matrix. More precisely, in the limit of a large number of neurons, the vector of tuning curve derivatives, **f′**, must be embedded in the subspace spanned by the first few eigenvectors of the covariance matrix. Importantly, for a finite number of neurons, we find in simulations that the more **f′** is embedded in the subspace, the lower the information. We take advantage of this observation to estimate information. We show with a model of V1 circuitry that this method can indeed detect differential correlations corresponding to discrimination thresholds of 1 degree, with experimentally attainable numbers of trials and neurons. We apply this method to recordings from awake monkeys, and find differential correlations corresponding to discrimination thresholds of 0.47-1.20 degrees, suggesting that sensory and behavioural information are comparable, and thus that neural circuits are efficient.

## Results

For simplicity, we consider a scalar stimulus, denoted *s*. What we want to know is how much information an area in the brain contains about that stimulus. As mentioned above, that would be easy if we could record from all (or a large fraction of) the neurons. But with current technology, we can’t; instead we have to extrapolate from a finite number. Unfortunately, naïve extrapolation turns out not to work well (Kohn et al. 2016). We can typically record from 100s of neurons, and in that range a plot of information versus number of neurons often does not exhibit clear signs of saturation.

We thus take a different approach, derived from our knowledge of the nature of neural noise in large populations of neurons. This approach starts with the usual tuning curve plus noise model: **r** = **f**(*s*) + noise where **r** is a vector of population responses and **f**(*s*) is a vector of tuning curves. (In components, *r*_*i*_ is the response of neuron *i* and *f*_*i*_(*s*) is the mean response of neuron *i* when stimulus *s* is shown.) Decoding the neural response corresponds to finding the stimulus, *s*, for which **r** is as close as possible to **f**(*s*). As can be seen in **FIG 2**, noise that has a component in the **f′** (*s*) direction is especially bad for decoding. In fact, decoding performance is determined almost solely by the amplitude of the noise along that direction. Such noise is known as differential correlations (Moreno-Bote et al. 2014).

**Figure 2.**
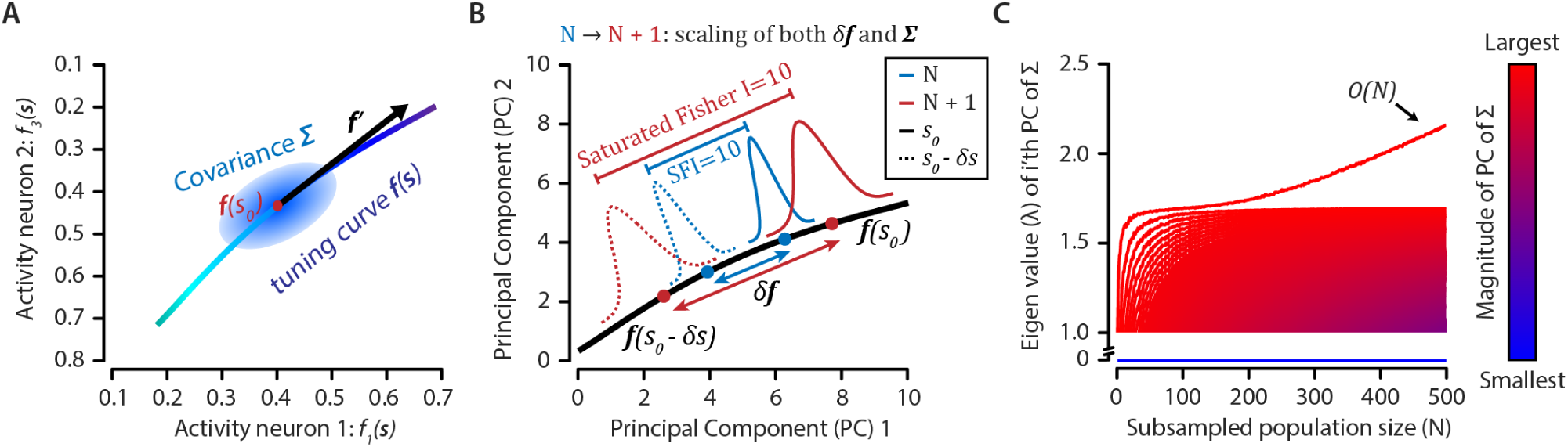
(**A**) Population patterns of neural activity can be thought as points in an *N* dimensional space in which each axis corresponds to the activity of a single neuron (out of *N* neurons in the population). As the value of the encoded variable varies smoothly, the population activity spans a one-dimensional nonlinear manifold whose shape is determined by the tuning curve, **f**(*s*), of the neurons. Variability in responses to a fixed stimulus *s*_*0*_ is represented as the blue ellipse centered at **f**(*s*_0_). (**B**) In a fine discrimination task, the ability to detect a small stimulus change, from *s*_*0*_ to *s*_*0*_*-δs*, depends on the distance between the average population activity for *s*_*0*_ and *s*_*0*_*-δs* and the amplitude of the neural variability. For *δs* small enough, the average activity manifold can be linearly approximated by a line that is tangent to the manifold given by **f**(*s*); this corresponds to the derivative of the tuning curves evaluated at *s*_*0*_, **f**′(*s*_*0*_). The projection of the neural variability along this axis is what is known as differential correlations. (**C**) Only a finite number of eigenvalues of the covariance matrix of the neural activity, **Σ**, can be O*(N)*. The covariance depicted here was obtained by injecting high orientation noise into our model (M1) (σ = 5°).

An important aspect of differential correlations is that when there is a finite amount of information in a population of neurons, the size of the differential correlations must be *O*(*N*); that is, the variance of fluctuations in the **f′** (*s*) direction must be proportional to the number of neurons, *N*, in the large *N* limit (Moreno-Bote et al. 2014). Consequently any eigenvector of the noise covariance matrix that has appreciable overlap with **f′** (*s*) must have an *O*(*N*) eigenvalue (**FIG 2c**). This suggests an alternate approach for determining the value at which information saturates, at least qualitatively: estimate the amplitude of differential correlations by enumerating the eigenvectors that have appreciable overlap with **f′** (*s*). The larger the fluctuations in the **f′** (*s*) directions, the more likely it is that **f′** (*s*) will strongly overlap with the eigenvectors with largest eigenvalues. Thus if differential correlations are strong (and information is low), the angle between **f′** (*s*) and the subspace spanned by the first few eigenvectors should be nearly zero (here and in what follows, we’ll rank eigenvectors by their eigenvalues). If, on the other hand, differential correlations are weak (and thus information is large), the angle between **f′** (*s*) and the subspace spanned by the first few eigenvectors should be large. In the next section we develop a quantitative measure based on these ideas.

### *φ*: a measure of differential correlations

Our approach to estimating the value at which information saturates is simple in principle: compute the angle between **f′** (*s*) and the subspace spanned by the first *k* eigenvectors of the noise covariance matrix (**FIG 3a**), denoted, cos^2^θ_*k*_, and plot cos^2^θ_*k*_ versus *k*. If differential correlations are strong this plot should go rapidly to 1; if differential correlations are weak this plot should go slowly to 1. To obtain a quantitative measure of “slow”, we plot the same thing, but for the shuffled covariance matrix. This gives us two curves – one for the true and one for the shuffled covariance matrix. We then define *φ* to be the area between them, normalized by *N* (see **FIG 3b**). In the presence of *O*(*N*) differential correlations (which, recall, is the case whenever there is a finite amount of information in the population) and in the limit of an infinite number of neurons and trials, *φ* should be close to 1/2. That’s because cos^2^θ_*k*_ is close to 1 when *k*/*N* is negligibly small (Blue curve, **FIG 3b**). For smaller populations, though, *k*/*N* is necessarily larger when cos^2^θ_*k*_ is close to 1 (simply because *N* is smaller). Therefore, the critical question is: what level of differential correlations can be detected reliably with the number of trials and neurons typically used in experiments?

**Figure 3.**
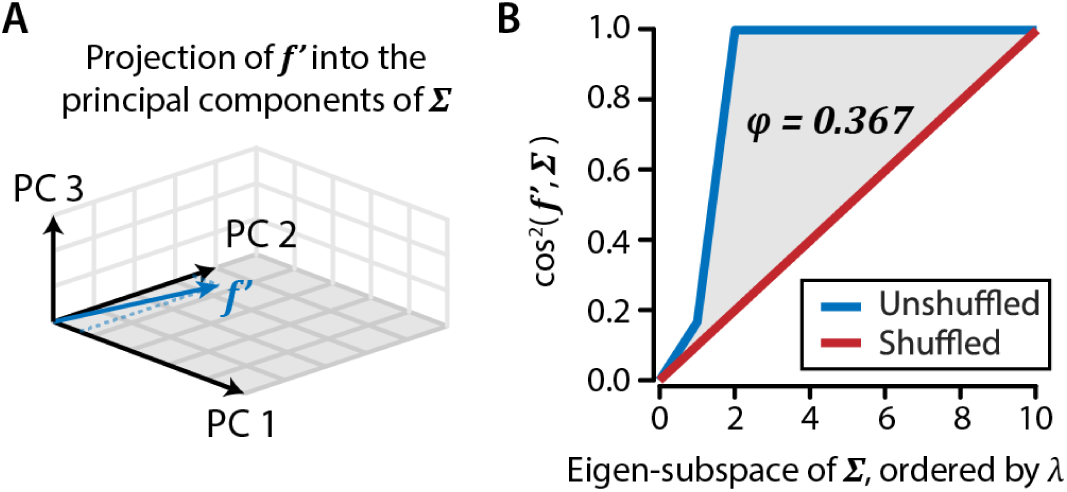
The alignment of the tuning curve’s derivative, **f′**, with the principal components of the covariance matrix can reveal differential correlations. (**A**) Schematic showing the calculation of the alignment of **f′** with the principal components (PCs) of the covariance matrix (***Σ***). (**B**) The difference in the area-under-the-curve (AUC) between the real (unshuffled, blue) and shuffled (red) alignment (cos^2^) of **f′** with the consecutive eigen-subspace of **Σ** provides a measure, denoted *φ*, for the strength of differential correlations.

We first explored this question through simulations. These simulations also allowed us to address one limitation of our technique, namely, that while the presence of strong differential correlations implies that our measure, *φ*, is large, the reverse is not true: *φ* can be large without *O*(*N*) differential correlations. As we will see, our simulations demonstrate that, in realistic simulation of cortical circuits, *φ* is small when differential correlations are small, and grows with the amplitude of the differential correlations.

## Simulations

To assess the sensitivity of ***φ*** to differential correlations, we used realistic simulations of cortical circuits, and computed ***φ*** as a function of the strength of the differential correlations, the number of neurons, and the number of trials. We performed simulations using two biologically plausible computational models of the early visual system. The first model (M1) used conductance-based leaky integrate-and-fire (LIF) model neurons for all cortical cells, and uncoupled point processes for pre-cortical stages (retina and LGN) (Somers, Nelson, and Sur 1995; Seriès, Latham, and Pouget 2004). ON-centre and OFF-centre inputs fed into V1, giving rise to Gabor-like receptive fields. V1 cells were also sparsely recurrently connected (**FIG 4**, see Methods for a more detailed description). The second model (M2) was a simplified version of the one presented in Kanitscheider et al. (2015). It was different from M1 in three respects: there were no lateral connections in the V1 layer, it used rate-based, rather than leaky integrate-and-fire, neurons, and differential correlations were introduced differently (see section “*φ* is proportional to the amplitude of differential correlations”).

**Figure 4.**
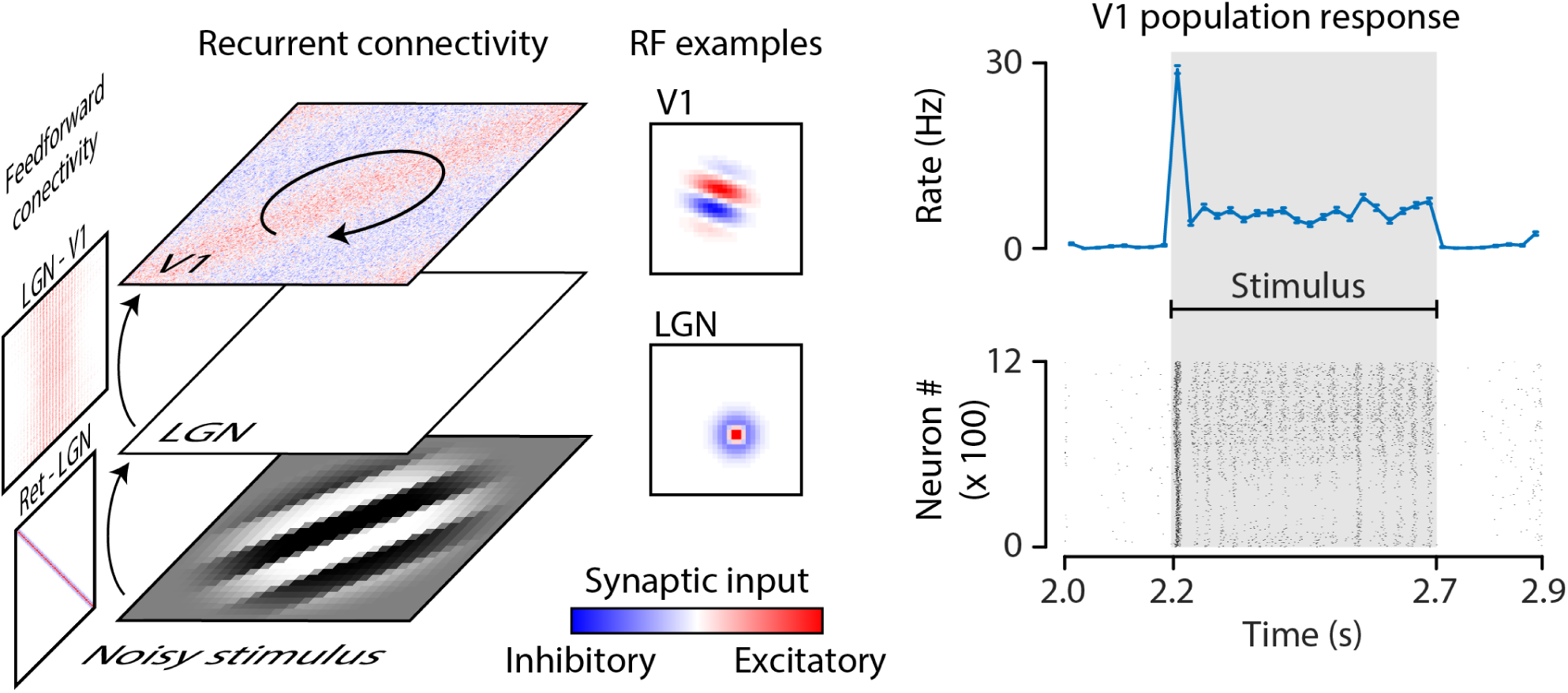
Summary of the computational model of the early visual system. We used biologically plausible feedforward connectivity between layers and recurrent connectivity within layers; that successfully generated orientation-tuned receptive fields in V1. An oriented grating was presented to the stimulus, and differential correlations were induced by introducing random orientation noise. The mean population response of all 1200 neurons to the stimulus (presentation epoch marked as Stimulus) is shown on the right as well as the spike raster over the same time period.

### *φ* is close to 0 in the absence of differential correlations

As mentioned above, when there are *O*(*N*) differential correlations and we can record from an infinite number of neurons, *φ* must be close to 1/2. When we can record from only a finite number of neurons, however, how *φ* scales with the strength of the differential correlations is unclear. At the very least, *φ* should go to zero as the strength of the differential correlations decreases. To test this prediction, we asked whether *φ* is significantly different from 0 in simulations in which differential correlations are *O*(1) by construction. To generate *O*(1) differential correlations, we simulated a variation of model M1 in which each cortical neuron received its own private set of LGN inputs, thus ensuring that the weighted sums of LGN inputs onto each neuron were statistically independent. While this ensured that no correlations were present in the LGN input arriving at V1, cortical neurons were still correlated due to the presence of lateral connectivity. This connectivity induced a low rank covariance matrix similar to what is observed *in vivo* (Huang et al. 2019) (**FIG 5)**. As can be seen in **FIG 6A**, in this case *φ* was close to zero (*φ* = 0.041). It is reassuring to see that *φ* is close to 0 in the absence of differential correlations in realistic simulations. This addresses the point we raised earlier: while, in theory, it is possible for *φ* to be large when differential correlations are small, this is not the case in our simulations.

**Figure 5.**
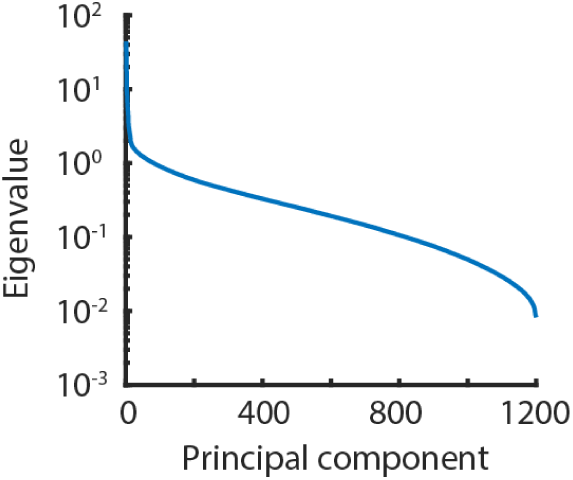
Example eigenspectrum of **Σ** in the absence of strong differential correlations. The sharp drop in magnitude of the eigenvalues (from ∼40 to ∼1) within the first couple of PCs is an indication that this covariance matrix is effectively low-rank.

**Figure 6.**
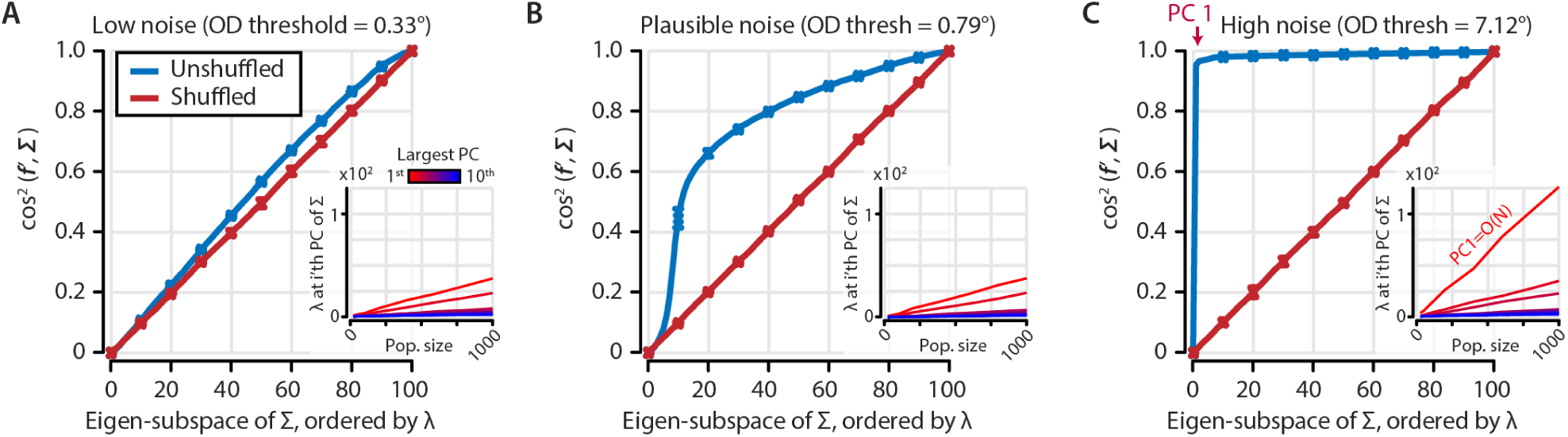
Differential correlations can be induced in the computational model with an injection of orientation noise, and show behaviour consistent with theoretical predictions. (**A**-**C**) Alignment of **f′** with the principal components of the covariance matrix **Σ** of a subpopulation of 100 simulated neurons for low (**A**), plausible (**B**), and high (**C**) levels of orientation noise, showing both unshuffled (blue) and shuffled (red) data. Insets show the growth of the 10 largest eigenvectors (principal components: PCs) of **Σ** as a function of population size. Note that with strong orientation noise, the first PC (PC1) is *O*(*N*); this generates the large jump in alignment with **f′** in panel C. The orientation discrimination thresholds (ODTs) above the graphs indicate the equivalent ODT according to the Fisher information calculated from the total population (N=1200 neurons). *φ* was 0.041, 0.373, and 0.481 for A-C respectively.

### *φ* is proportional to the amplitude of differential correlations

Next, we asked whether *φ* increases with the strength of the differential correlations, using simulations of model M1 with LGN inputs that were shared among V1 neurons. To modulate the strength of differential correlations, we injected noise into the orientation of the gratings presented to the network. We did this by presenting gratings at slightly different orientations on each trial. In our simulations, the orientations were drawn from a Gaussian distribution with standard deviation σ_θ_, which is proportional to the orientation discrimination threshold (eqs 16-18). As expected, *φ* grew monotonically with the orientation noise, σ_θ_, and thus monotonically with the strength of the differential correlations (**FIG 6, B-C**. We found that with 100 neurons and 805 trials (the average number of neurons and trials across all experimental recordings), *φ* becomes statistically different from zero when the orientation noise was around 0.4 degrees, corresponding to a discrimination threshold of 0.59 deg in V1 (**FIG 7A**).

**Figure 7.**
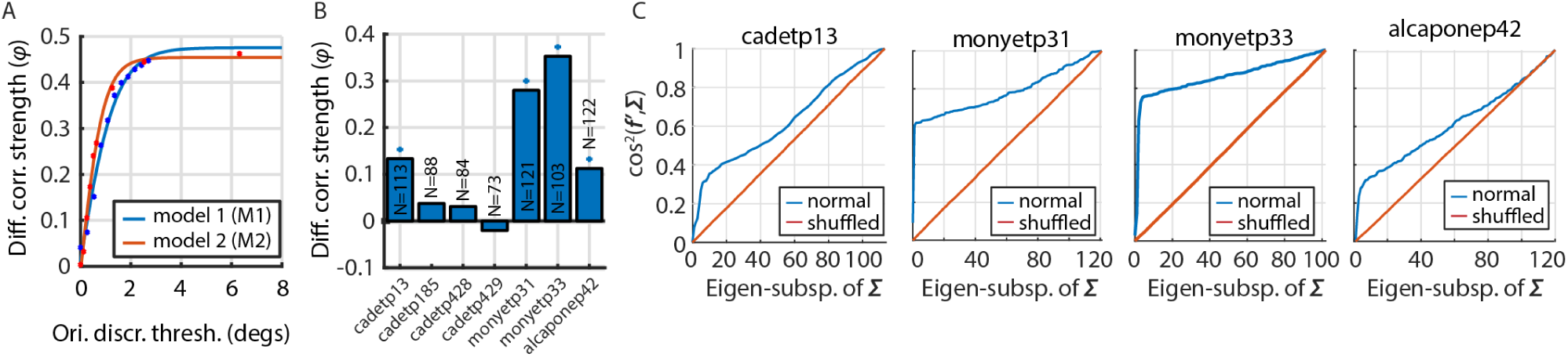
Differential correlations in experimental data. (**A**) A series of simulations with two models (blue, red) and varying orientation noise injections (dots). This was fit with a logistic sigmoid (solid curves), which allowed us to estimate the equivalent orientation discrimination threshold for the mean *φ* value observed in the experimental data. (**B**) Data from 4 out of 7 data sets (blue +) recorded in V1 show a signature of information-limiting correlations (see C). (**C**) *φ* curves for the four data sets that showed a significant effect (p=7.0 ×10^−5^, p=3.9 ×10^−15^, p<1.0 ×10^−308^, and p=1.6 ×10^−3^ respectively).

For model M2, we injected noise in the response of the retinal cells, as opposed to the orientation of the grating. As shown in Kanitscheider et al (2015), this retinal noise, when large enough, generates large differential correlations and leads to strong information saturation. As in M1, we found that *φ* grew monotonically with the amplitude of the injected noise. When plotted as a function of the discrimination threshold of the V1 layer, the resulting curve is very similar to the one we obtained with model M1 (**FIG 7A**). These second simulations show that the *φ* measure is not overly sensitive to the details of the model and injected noise.

### Experimental data

Having verified the validity of our methods *in silico*, at least for a set of reasonably biologically plausible models, we analysed 7 data sets recorded in the primary visual cortex of three awake monkeys (**FIG 7**). In all cases the data sets came from chronically implanted Utah arrays. Data sets 1-4 came from monkey 1; data sets 5 and 6 from monkey 2; and data set 7 from monkey 3. In every animal, *φ* was significantly different from zero, though this was not the case in all sessions in monkey 1 (**FIG 7B**). Therefore, in all three monkeys, we found evidence of differential correlations in V1.

To relate the level of differential correlations present in these data sets to the information the V1 population would be expected to provide for orientation, we converted *φ* into discrimination threshold (eqs 16-18) using the plot shown in **FIG 7A**. This analysis returned a range of thresholds from 0.47 to 1.20 degrees (0.51, 0.87, 1.20 and 0.47 degrees for, respectively, data set 1, 5, 6 and 7).

## Discussion

We have presented a new method that can detect the presence of differential correlations corresponding to discrimination thresholds of 0.47 to 1.20 degrees in the spiking activity of 100 neurons and <1000 trials. Such a method is useful because the vast majority of simultaneous recordings of spiking activity are currently limited to a few 100 neurons and information saturation in populations of this size may not always be evident.

When applied to V1 recordings in awake monkeys, our approach revealed, in four data sets, the presence of strong, information limiting, differential correlations. However, *φ* was not significantly different from zero in the remaining three data sets. Intriguingly, these three data sets were obtained from the same implant as a data set in which the value of *φ* suggested a discrimination threshold of 0.51 degrees, several months earlier. Assuming the representational quality of V1 was not altered by the implant, the reduction in the value of *φ* might reflect the established degradation of recording quality in chronic arrays over time (Barrese, Aceros, and Donoghue 2016).

In the data sets for which *φ* was statistically significant, the corresponding discrimination thresholds were on the order of 1 degree. This is typical of the behavioural thresholds that have been reported experimentally (∼1 deg, (Webster, Switkes, and De Valois 1990)). This argues that information does not saturate at extremely high levels, as suggested by several groups (Shamir and Sompolinsky 2006; Ecker et al. 2011) (illustrated in **Fig 1**). In contrast, it is consistent with a previous report in V1 which found discrimination thresholds of the same order of magnitude as the behavioural one (2.7% vs 4.8% contrast detection threshold) (Chen, Geisler, and Seidemann 2006). The similarity we found in V1 between behavioural and neural thresholds has been reported in MSTd, where it was found that about 80% of the information in that area is used to drive the behaviour of animals engaged in a heading discrimination task (Pitkow et al. 2015; Kim et al. 2016).

A recent rodent study using calcium imaging of more than 20,000 neurons also demonstrated the presence of strong differential correlations in V1 (Stringer, Michaelos, and Pachitariu 2019). The authors found that information saturates as the number of neurons goes to infinity, and the discrimination threshold at saturation is about 0.3 degrees. This is smaller than typical discrimination thresholds in mice (which are about 5 degrees; (Glickfeld, Histed, and Maunsell 2013)). However, the 0.3 deg discrimination threshold was derived from activity presented in a full hemifield stimuli for 750 ms. It is unlikely that mice can integrate information over the entire visual hemifields and over such a long time. Once these constraints are taken into consideration, the gap between behavioural and neural threshold will almost certainly shrink.

Altogether, these results suggest that neural circuits downstream of V1 of macaque monkeys are not vastly inefficient, in the sense that information in neural responses is about the same as information in behavior. Nonetheless, information loss occurs. This could be the result of internal noise in neural activity, but unless the noise is specifically aligned with the **f′** (*s*) direction and is O(*N*), internal noise cannot significantly affect information transmission (Zylberberg et al. 2017). A more likely source of information loss is suboptimal computation (Beck et al. 2012). For instance, and as mentioned earlier, animals may not integrate information properly across space and time, and may not use the optimal synaptic weights between cortical areas. This will necessarily lead to information loss in downstream areas which will be reflected in a reduction in the norm of **f′** (*s*), an increase in differential correlations, or a combination of the two.

## Methods

### Computational model for large-scale simulations

#### Summary of network architecture

Model M1 contains two types of model neurons, which are described in more detail below. Pre-cortical stages, i.e. Retina and LGN, are uncoupled point processes, where the Retina is a filter-based 2D activity map with i.i.d. Gaussian noise on top of the filter response, and the LGN turns this activity map into spikes, using a stochastic firing probability. This LGN stage feeds into the first fully simulated cortical stage (V1), using a connectivity pattern that gives rise to Gabor-like receptive fields in V1, similar to (Seriès, Latham, and Pouget 2004) and (Somers, Nelson, and Sur 1995). All cortical cells are simulated using a conductance-based leaky-integrate-and-fire model. V1 cells are sparsely recurrently connected, as described below.

#### Filter-based input layers: Retina, LGN

The Retina layer in our model consists of two separate streams (ON-centre and OFF-centre) that are arranged in arrays of 32 × 32, with a spacing of 0.2 degrees visual angle between cells. For a patch of 6.4 × 6.4 degrees we simulated a visual input twice the size (12.8 × 12.8) to avoid edge effects. The rate-based activation in Retina for a cell at location (*x,y*) for ON-centre and OFF-centre streams were:

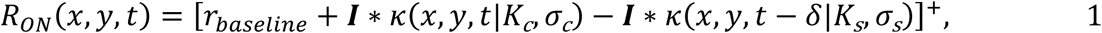

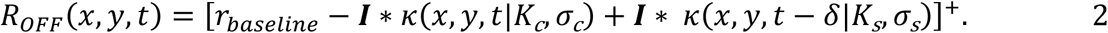

Here, [·]^+^ =max(·,0) denotes rectification, ***I*** is the input image, c indicates centre response, s indicates surround response and δ is a 3 ms delay between centre and surround responses. These responses are in turn generated by convolving the visual stimulus ***I*** with a spatiotemporal kernel *κ*, consisting of a circularly symmetric Gaussian spatial profile *G* and an exponential temporal impulse function *F*:

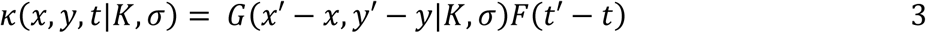

Here, the *x’, y’* and *t’* denote the spatiotemporal location of the filter in the corresponding *x, y, t* space of the input image ***I***. We truncated the filter for locations more than four times the standard deviation of the Gaussian profile away from its mean for computational efficiency. The spatial circularly symmetric Gaussian *G* and decaying exponential function *F* are defined as:

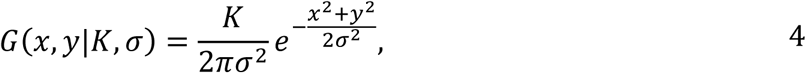

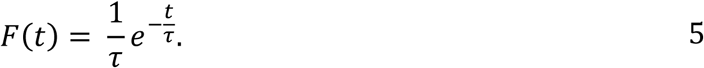

We used the following parameters, taken from (Somers, Nelson, and Sur 1995): *σ*_*c*_ = 0.176, *σ*_*sd*_ = 0.53, *K*_*c*_ = 17, *K*_*s*_ = 16, *τ*_*c*_ = 10, *τ*_*s*_ = 20, and *r*_*baseline*_ = 15. The LGN layer simply transforms the R_ON_ and R_OFF_ analogue activity levels into stochastic spiking events, with an average rate of 40 Hz at full stimulus contrast.

#### LGN-V1 connectivity structure: Gabor fields

Each V1 cell has a Gabor-like receptive field, with several preferred parameters that varied across cells: phase (ψ), orientation (θ), spatial frequency (1/λ), spatial length (σ_y_), spatial width (σ_x_), horizontal centre (ξ), and vertical centre (υ). We created a Gabor patch G by combining these parameters as follows:

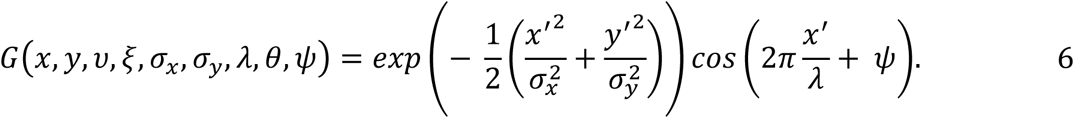

Here *x’* and *y’* are defined as:

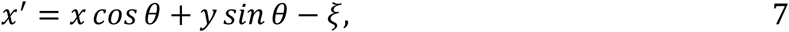

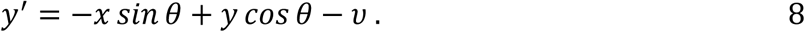

With each V1 cell assigned a Gabor field, we next calculated the connection probability for ON-centre cells at retinoptic locations (*x, y*) from LGN to a V1 cell as follows,

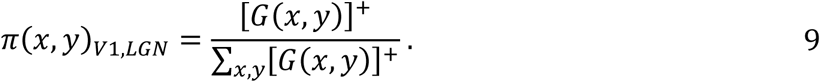

For OFF-centre cells, we used the same formula, but inverted the sign of *G*. We then connected the required number of LGN cells (72 for pyramidal cells, 48 for interneurons) from each stream to a V1 cell by choosing an LGN cell to project with a probability proportional to its weight described above, relative to that of other cells. If a connection was made, we assigned to each a weight equal to its corresponding probability, *π(x,y)*, a conductance (0.264 and 0.288 for pyramidal cells (E) and interneurons (I) respectively) and a Gaussian delay (E: 0.01 ± 0.007, I: 0.05 ± 0.003; mean ± SD, rectified to ≥ 0).

#### Leaky-integrate-and-fire (LIF) layer V1

For the fully simulated cortical layers, we used a conductance-based leaky-integrate-and-fire model of pyramidal cells and interneurons, as described in (Seriès, Latham, and Pouget 2004) and originally in (Somers, Nelson, and Sur 1995). We simulated 1200 V1 cells, of which 80% were excitatory and 20% were inhibitory. For post-synaptic cell *i*, the membrane voltage, *V*_*i*_, evolves according to

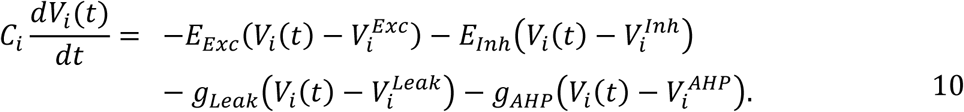

Here, *E*_*Exc*_ denotes the total excitatory post-synaptic potential (PSP) due to feedforward and recurrent connections, *E*_*Inh*_ denotes the total inhibitory PSP, *g*_*Leak*_ denotes the passive leak conductance, and *g*_*AHP*_ the after-hyperpolarization conductance due to prior spiking (see eqs. 11-13 for the calculation of the PSPs). *V*^*Exc*^, *V*^*Inh*^, *V*^*Leak*^ and *V*^*AHP*^ denote the reversal potentials for excitatory, inhibitory, leak and AHP inputs respectively. *V*_*i*_*(t-dt)* denotes the cell’s membrane voltage at the previous time step, and *C*_*i*_ the membrane capacitance for neuron *i*. Whenever a cell’s membrane voltage exceeded −55 mV, we used a delta function to simulate spiking behaviour, and reset the cell’s membrane voltage to its resting potential (*V*^*Leak*^) on the next time step. As the simulation time step was 0.5 ms, our cells had an absolute refractory period of 1 ms. We used the following (approximately) biologically plausible parameters for all cells: *V*^*Exc*^ = 0 mV, *V*^*Inh*^ = −70 mV, *V*^*Leak*^ = −65 mV, *V*^*AHP*^ = −90 mV. For excitatory cells, we used *g*_*Leak*_ = 25 nS, *g*_*AHP*_ = 40 nS, *C*_*i*_ = 0.5 nF; and for inhibitory cells we used *g*_*Leak*_ = 20 nS, *g*_*AHP*_ = 20 nS, *C*_*i*_ = 0.2 nF. The membrane voltage at t = 0 was chosen randomly according to a Gaussian distribution with μ = −55.6 and σ = 1.

We increased computational efficiency of our model by vectorising the calculation of spike-induced post-synaptic potentials across spike history and synaptic connections, thereby requiring at each time step only a for-loop across cells, rather than across synaptic connections. This is difficult to achieve in a setup such as ours, where heterogeneous synaptic delays and exponentially decaying PSPs act on a single target neuron, because calculating the resulting action on the membrane voltage requires incorporating all heterogeneous pre-synaptic spike times from various time steps in the past. We achieved this vectorization by using a sparse dynamic cylindrical array to store recent spiking activity for all neurons in a reusable buffer variable, as described originally by (Brette and Goodman 2011). The algorithm (see below for a pseudo-code description) uses a vector ***ν*** that stores the number of recently emitted spikes per cell, a vector **π** that keeps track of the current position on the cylindrical array for each cell, and a cylindrical array **T** of size [cells x spikes] that stores the time stamps of recent spikes. Looping across cells, single-synapse PSPs are stored in a vector, where each element *q* corresponds to a single synaptic connection from cell *j* to cell *i*. For each source cell *j*, the PSPs in all its output synapses can be calculated using vector operations. We loop through all neurons in ***ν***, ignoring all cells that have zero recent spikes, yielding another performance boost. Let *n* be a non-zero element for cell *j* in **ν**, and **p** be an *n*-dimensional vector of indices computed from ***π*** that correspond to the locations in **T** where the most recent spikes are stored. Now, the recent spike times of cell *j* can be accessed simply through **T**(*j*,**p**), providing a vector ***τ*** of time stamps. We next created a matrix **Δ** of size [spikes x synapses] that contains the spiking time relative to the current simulation time *t*, normalised for the synaptic delays **δ**. For spike *k* and synapse *l*, this relative spiking time can thus be written as:

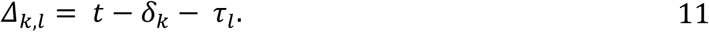

Note that this operation is easily written as a vectorised calculation in any Matlab implementation. The PSPs ***s*** for all target synapses of *j* are now given by:

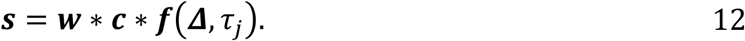

Here, * indicates element-wise multiplication, ***w*** is a vector of synaptic weights, and ***c*** is a vector of synaptic conductances, *τ*_*j*_ denotes the synaptic time constant for cell *j*, which depends only on whether the cell is excitatory or inhibitory, and ***f*** is the exponential PSP function that outputs a sum total value for each synapse. As above, note that this function is completely vectorised over synapses, but for clarity we here write the function for a single synapse *l*:

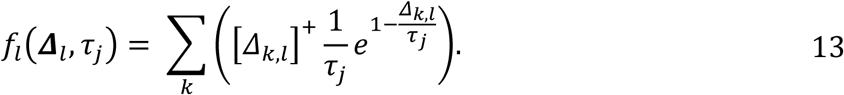

We used the following parameters: *τ*_*Inh*_ = 2 ms, *τ*_*Exc*_ = 1 ms, and cylindric buffer size *t*_*buffer*_ = 200 ms. Synaptic delay and conductance parameters are provided in their corresponding paragraphs detailing the connectivity structure. Finally, the resulting excitatory and inhibitory input per cell (*E*_*Exc*_ and *E*_*Inh*_) can be calculated by summing all synaptic PSPs in **s** to their postsynaptic targets. This step is also easily vectorised, using the built-in Matlab function *accumarray()*. In Matlab pseudo-code, this algorithm can be written as follows:

Matlab pseudo-code description for the algorithm based on (Brette and Goodman 2011)

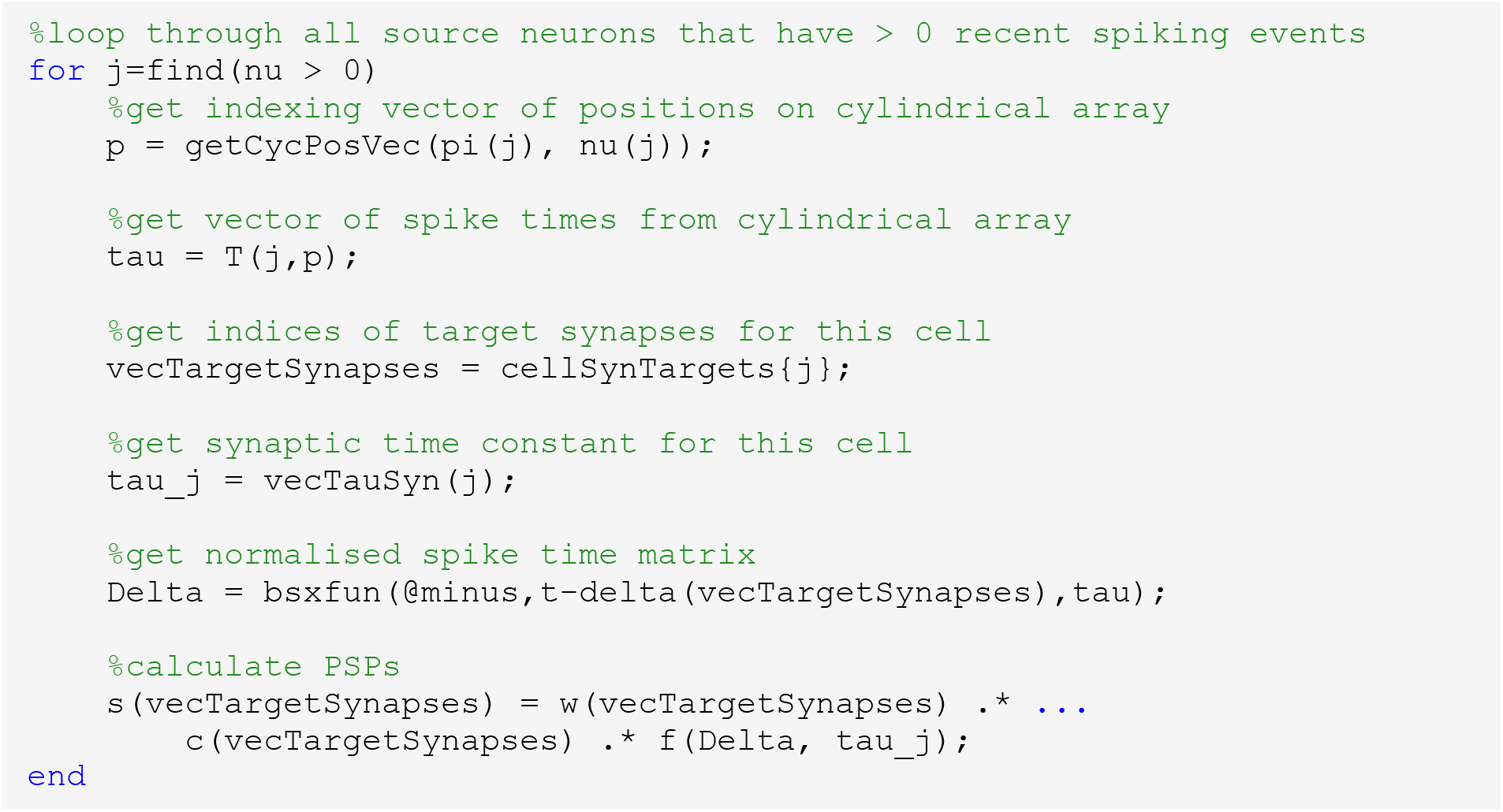

#### V1 recurrent connectivity: similarity-& locality-based

Our V1 connectivity was determined on the basis of receptive-field similarity, where the weighting of the similarity is controlled by a locality hyperparameter *λ*, in the interval [-1 1]. In this range, λ = −1 corresponds to a uniform connectivity (i.e., ignoring similarity), λ = 1 corresponds to a maximally local connectivity (i.e., connect *n* most similar cells), and λ = 0 corresponds to a proportional weighting, as described above for LGN-V1 connectivity. We set λ = 0 for excitatory neurons, and λ = −0.5 for inhibitory neurons. This means that inhibitory neurons project more uniformly to less similarly tuned cells than excitatory neurons, as has been shown in mouse visual cortex (Ko et al. 2011). We calculated an element-wise similarity *ρ* in receptive fields of cells *i,j* as follows. First, we normalised each receptive field **F** of size [*x* by *y*] to be mean-zero:

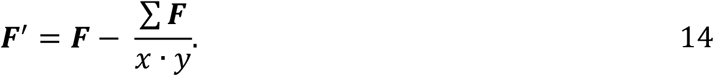

Here, *Σ* **F** denotes the sum over all elements in **F**, and · denotes a scalar multiplication. The similarity metric *ρ* (in the range [-1 1]) is given by

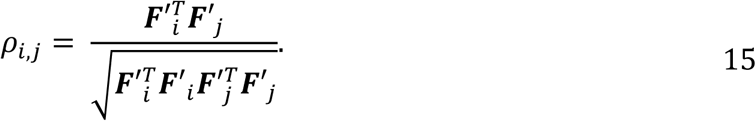

Here, ^T^ indicates transpose. We rectified all negative *ρ* values to 0, and created a cumulative probability vector **l** based on [*ρ*]^+^, as described above for LGN-V1 connectivity. For a number of connections *n* originating from a source neuron, we randomly chose *n*_*proportional*_ *= n\**(*1-*|*λ*|), rounded to the nearest integer, connections based on **l**. The remaining *n*_*non-proportional*_ *= n* -*n*_*proportional*_ connections were chosen either according to a uniform random distribution across neurons when *λ < 0*, or the *n*_*non-proportional*_ most similar cells were connected when *λ > 0*. This way, connectivity is a mixture of uniform and [*ρ*]^+^-proportional connections for -*1 < λ < 0*, and a mixture of most-similar and [*ρ*]^+^-proportional connections for 0 *< λ < 1*.

#### Programmatic implementation

The computational model was programmed in Matlab R2015b, compiled as a stand-alone executable, and run in a massively parallel manner on the University of Geneva High Performance Computing cluster *Baobab*. Each cluster node ran a set of stimulus repetitions with randomised initialization parameters, and the resulting simulated data were post-hoc combined into a single data set.

#### Visual stimulus parameters

Visual stimuli were always centred in the middle of the simulated patch of visual space and consisted of drifting sinusoidal gratings within a circular aperture 5° in diameter, with a cosine-ramped edge (period of 1°) leading to a neutral-grey background. Unless stated otherwise, all stimuli had a pre-stimulus uniform neutral-grey blank period of 100 ms, after which the stimulus was presented for 500 ms, and ended with another 100 ms blank period. We used a spatial frequency of 0.25 cycles per degree, a temporal frequency of 2 Hz, full contrast and luminance, and a random starting phase. We presented a single pair of orientations separated by five degrees (42.5 and 47.5 degrees from vertical).

We can quantify the amount of information encoded about a stimulus at both the neural level as well as at the final behavioural level. A comparison of these two quantities would tell us whether neural computations are efficient or not. If information about the stimulus in V1 exceeds the information we can extract from the behavioural output, then computations must not have been efficient; the difference must have been lost as information was transmitted up the cortical hierarchy to frontal regions and ultimately to the behavioural motor output. Conversely, if V1 information saturates close to behavioural information, then all intermediary neural computations between the two levels must have been extremely efficient.

Behavioural information is usually quantified using the just-noticeable-difference (JND), defined as the smallest orientation difference where the subject is still able to report accurately which of two orientations is more tilted clockwise. For humans, this is around 1° (Webster, Switkes, and De Valois 1990), with the JND being defined at the orientation where responses were on average 75% correct.

Neural information is usually quantified instead using Fisher information, and in the rest of this paper, we shall use this metric rather than the JND; compared to a JND analysis, it has some desirable features such as its applicability to neuronal activity within the framework of information-limiting correlations (Moreno-Bote et al. 2014). Thus, to compare information at the two levels, we must convert the JND into its equivalent Fisher information.

For a scalar stimulus, linear Fisher information (defined in a later section) can be related to the sensitivity index *d’* by (Averbeck and Lee 2006; Seung and Sompolinsky 1993)

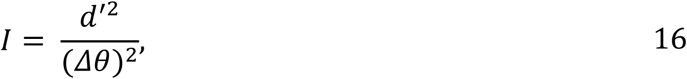

and in visual space, *d’* is defined by

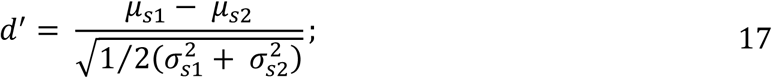

here, *μ*_*s1*_ and *μ*_*s2*_ are the means of the two noisy stimuli and 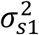 and 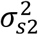 their variances. JND is measured using a 1AFC task, and for such a task, the proportion-correct can be related to *d’* by (Micheyl, Kaernbach, and Demany 2008)

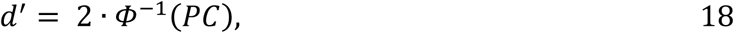

where *Φ*^*-1*^*(x)* is the inverse of a cumulative Gaussian from *-*∞ to *+x*. Thus using (17), we can convert a 75% proportion-correct into the *d’* value of 1.35, and using (16), we can convert this into a Fisher information of *I* = 1.82 for an orientation discrimination threshold of 1 degree.

It is currently not possible to directly measure the information in V1, as the required number of trials and neurons still vastly exceeds the capabilities of even the latest recording techniques. Therefore, we instead used a biologically plausible computational model of early visual primate cortex to simulate the strength of information-limiting neural correlations when orientation information saturates at a 0.85 deg orientation discrimination threshold; that is, our model simulates neural activity assuming information would not be lost in the subsequent neural computations up to the motor output. To do this simulation, we artificially saturated the input information in our model at the required level by randomly perturbing the stimulus with orientation noise. In effect, we created a non-zero variance in the feature dimension over multiple presentations of the same stimulus. Using equation (17) with 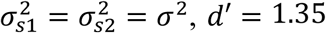, and Δ*μ* = 0.85°, we deduced the necessary orientation-noise level to be around σ = 0.63°. Using equation (17) with Δμ = 2° instead for a 2-degree discrimination threshold, the necessary orientation-noise level is σ = 1.48°.

### Analyses for simulated and experimental data

#### Data pre-processing and general steps

We transformed all spike-time based data into a matrix **A** of total spiking activity during stimulus presentation per neuron and trial, giving a [*t* x *m*] spike count matrix of *m* neurons and *t* trials. Unless stated otherwise, our shuffling procedure shuffled neural responses per neuron across repetitions of the same stimulus type. This shuffling procedure destroys all temporal correlations, but keeps single neuron firing statistics (e.g., tuning curves) intact.

#### Computations of Fisher information and f′

Much of our analysis involved estimating from data derivatives **f**′ of the tuning curve vector **f** with respect to the scalar stimulus parameter *θ*. This was done by finite differencing where

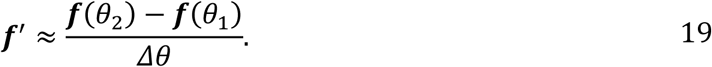

Linear Fisher information is defined to be

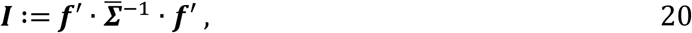

where 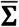 is the mean of the covariance matrices at *θ*_1_ and *θ*_2_, that is,

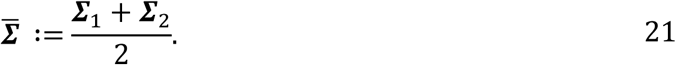

Linear Fisher information provides a lower bound on the total Fisher information encoded in the population. We note that in the 1-dimensional case, this definition of Fisher information (20) reduces to (16).

An estimator of 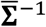 obtained by taking the matrix inverse of an estimator of 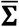 will generally be biased. We have therefore used a bias-corrected estimator of Fisher information as derived by (Kanitscheider et al. 2015) and given by

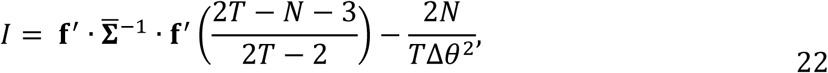

where *T* is the number of trials and *N* is the number of degrees of freedom. This estimator is only valid when *T* > (*N* + 2)/2. Where this condition is not satisfied, we forgo the bias-correction.

#### Projections of f′

Our analysis examined the behaviour of 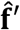 when projected into a variety of subspaces, where 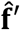 denotes **f**′ normalised. For all subspaces, the method of projection was the same. Suppose we are working in an *n*-dimensional neural activity space and are interested in projecting into a *k*-dimensional subspace. Then given a set of *n*-dimensional normal bases vectors {*v*_*i*_} spanning our subspace, the projection of **f**′ is given by

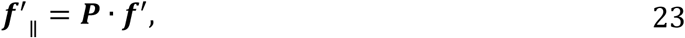

where **P** is the projection matrix given by

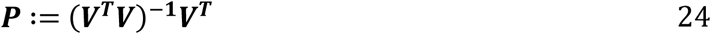

and **V** is the *n* × *k* matrix with columns corresponding to ***v***_*i*_; the projection of 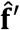 is simply

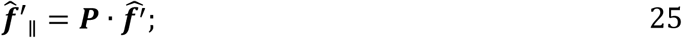

The projection of 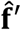 into 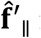 is quantified by the cosine between the two vectors; it can be shown that this is equivalent to

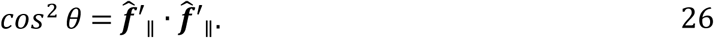

#### Bases vectors for the principal components of Σ

One of the subspaces considered was that spanned by the first *k* principal components of **Σ**. In this case, the bases vectors are orthonormal, and the projection matrix simplifies to

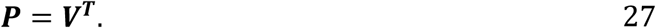

Moreover, it can be shown that the projection of **Σ** can be expressed as

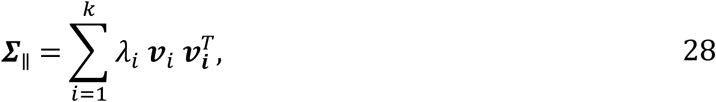

The cosine between 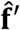 and 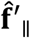 can be similarly simplified as well. Let 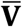 denote an *n* × *n* matrix where the columns comprise all *n* normalised eigenvectors of **Σ** arranged in order of descending eigenvalue, and let 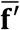 denote the vector given by

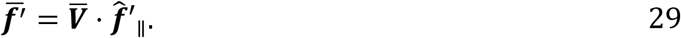

Then the cosine can be expressed as

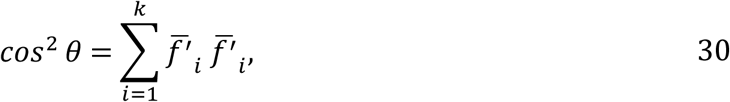

where 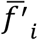 denotes the *i*th vector element of 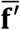.

#### Metric for differential correlation strength

We wanted to investigate whether experimentally recorded neural data shows differential correlations of a similar strength to what we expected from our computational model, where we limited the input information to levels that were behaviourally plausible (see the section on “Visual stimulus parameters”). To compare differential correlation strengths in data sets of varying quality and population sizes, we worked in terms of the fraction, *η*_*k*_, of the total resultant length (i.e., *cos*^*2*^*θ*) of 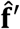 after projection into 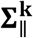, which we define as the *k*-dimensional subspace of **Σ** spanned by its first *k* principal components, using equations (28) and (30):

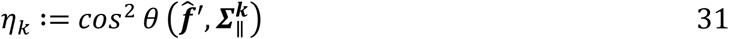

To quantify the strength of differential correlations, we defined a metric comparing the *cos*^*2*^*θ* in the real data set, 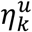, against that in a data set obtained after correlations are destroyed by shuffling, 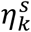; this metric is defined by

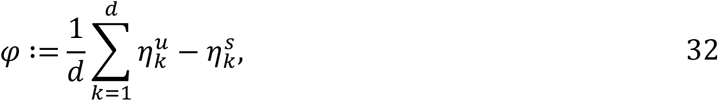

with *k* proceeding in order of the principal components of the *d*-dimensional covariance matrix **Σ**. This metric is 0 when the shuffled and unshuffled spaces have the same proportion of their respective total information across all subspaces. It is 1 when the unshuffled information in the 1-dimensional subspace is already equal to the total information, and the shuffled information is zero until the last dimension inclusive. *φ* is 0.5 in the case where shuffled information grows linearly, and unshuffled information is already equal to the total information in the 1-dimensional subspace. This latter case should be taken as a more realistic maximum value that *φ* could take.

#### Neurophysiology

General methods can be found in (Jasper, Tanabe, and Kohn 2019). In brief, animals were trained to fixate a small spot (0.2 × 0.2 deg), and maintain eye position within a 1-1.6 degree window. Eye position was monitored with a high-speed video tracking system (Eyelink II). After training, we implanted multi-electrode (‘Utah’) arrays (Blackrock; 1 mm length, 400 micron inter-electrode spacing). In two animals (Cadet and Monyet), we recorded V1 activity using one 96-channel and one 48-channel array. In the third animal, we implanted two 96-channels in V1. All animals were also implanted with arrays in area V4, not analysed here. Signals from each electrode were amplified and band-pass filtered (0.3-7500 Hz) using commercial acquisition systems (Blackrock Microsystems and Grapevine, Ripple), and sorted offline into single units and small multiunit clusters.

Visual responses were measured using full contrast, drifting sinusoidal gratings, presented on a calibrated monitor placed 64 cm from the animal (1024 × 768 resolution, 100 Hz refresh). After fixation was established and an additional delay of 100 ms, we presented a sequence of three gratings, each for 200 ms and followed by an inter-stimulus interval of 150 ms, during which a gray screen was presented. Responses were measured as the spike count in the 200 ms epoch during a stimulus was on the screen. Each recording involved either 2 gratings (1 session, orientation of the two gratings was 5 degrees apart) or 4 gratings (remaining sessions; consisting of two sets of gratings, with an orientation difference of 5 deg within a set, and 90 degrees between sets). All gratings had a spatial frequency of 2 cyc/deg, and a drift rate of 5 Hz. Grating size was between 2.5 and 8.6 deg in diameter. Animals worked an average of 913+/-206 trials, resulting in >2700 stimulus presentations per session.

In one animal (Cadet), we detected cross-talk between channels of the array (1-3% of pairings). Cross-talk was evident as frequent, precise synchronous activity between different electrodes. To address this issue, we removed, randomly, the spikes from one of the participating units, whenever a synchronous event occurred. In the remaining two animals, cross-talk was extremely rare (<0.1% of pairings), with the exception of three units in one animal (alcapone), which we excluded from our analysis.

